# Economic load-carrying in cockroaches

**DOI:** 10.1101/2025.09.23.678102

**Authors:** Be Eldash, Rudolf Johannes Schilder

## Abstract

Energetic costs of carrying loads can significantly impact animal fitness but appear to vary dramatically among animals. For some they equal the cost of carrying an equivalent amount of extra body mass while others carry loads more economically. Locomotor systems can plastically respond to acute and chronic loading, but how such responses impact energetics of locomotion is unclear. We asked how loading affects the energetics of *Blaberus discoidalis* cockroaches at rest and during locomotion at various speeds, and if energetics change as animals adjust to chronic loading. Cockroaches carried loads economically as early as 24 hours after load addition, with no change in energetic costs during a 10-day period. We discuss the implications of these findings, and potential mechanisms underlying economic load-carrying in arthropods.

## Introduction

Whether it is due to growth, seasonal oscillations in body mass or the need to carry resources, offspring and ornaments, all animals experience variations in load during their lifetime. The energetic cost of carrying a load can significantly impact an animal’s reproductive potential, even when costs represent a small fraction of the maximum metabolic rate (Berke et al., 2006). Thus, mechanisms that allow laden animals to minimize the energetic cost of carrying loads could be beneficial (Alexander, 1989; Berke et al., 2006). The systems that control and execute movements during locomotion are highly plastic and undergo adjustments in response to loading (Baldwin & Haddad, 2002; Turner, 2007; Schilder, 2016; Fox et al., 2020), but how such adjustments affect the energetic cost of carrying loads is unclear. This is relevant to our understanding of the costs associated with activities that are essential for survival and reproductive success, and of conditions in which plasticity fails to accommodate changes in load (Schilder et al., 2011; Bollinger, 2017; Fox et al., 2020).

In vertebrates, the energetic cost of carrying loads during legged locomotion equals that of moving an equivalent amount of actual body mass (Taylor et al., 1980). In other words, carrying a load consisting of half of one’s body mass results in a 50% increase in the cost of locomotion; carrying the equivalent to one’s body mass results in a two-fold increase, etc., irrespective of speed (Kram & Taylor, 1990). There are noteworthy exceptions: Women from the Luo and Kikuyu tribes (Maloiy et al.,1986), Nepalese porters (Bastien et al., 2005) and guinea fowl (Marsh et al., 2006; Katugam-Dechene, 2023) carry loads economically. For these groups, carrying a unit mass of load costs less energy than a unit of body mass. This is only true when subjects are habituated to carrying loads, suggesting that plastic responses to chronic loading may increase the economy of locomotion. Indeed, the study by Taylor et al. (1980) mentions that training led to an increase in the economy of locomotion of laden mammals, although no data were presented that supported this. Thus, there is some support for the hypothesis that adjustments to a chronic increase in load leads to an increase in economy of carrying loads, although it remains uncertain whether this is a general phenomenon (Sparrow & Newell, 1998).

Little is known about the energetics of chronic loading in invertebrates. For some species of ants, like vertebrates, the cost of locomotion is directly proportional to the total mass, irrespective of the distribution between body and load mass (Nielsen et al.,1982; Lighton et al., 1987; Bartholomew et al., 1988; Duncan & Lighton, 1994; Nielsen, 2001). Other invertebrates carry loads economically. Economical load-carrying was reported in cockroaches (Lee, 1994; Full et al., 1984 cited in Kram, 1996), crabs (Herreid & Full, 1986), and ants (Nielsen & Baroni-Urbani, 1990; Schilman & Roces, 2005). In an extreme example, rhinoceros beetles carrying loads equivalent to 10 times their body mass showed a less than two-fold increase in energetic cost of locomotion (Kram, 1996). Nonetheless, as in the vertebrate studies, most of these experiments examined acute effects of loading in animals that were accustomed to carrying loads (i.e., worker ants). We are unaware of studies on invertebrates that examined changes in energetics that occur as animals adjust to chronic loading.

In summary, carrying a load results in an increase in the energetic cost of locomotion that can have important consequences for chronically laden animals. Adjustments to load are known to affect locomotor performance, and there is evidence, at least in vertebrates, that they may impact the economy of carrying loads. Yet, this hypothesis has not been tested in invertebrates. We asked how loading affects the energetics of the cockroach *Blaberus discoidalis* at rest and during locomotor activity, and whether these costs change as cockroaches adjust to chronic loading. We hypothesized that 1) laden cockroaches will use energy at comparable rates to unladen controls at rest; 2) laden cockroaches will use more energy than unladen controls during locomotion, although this increase may not be as high as expected if carrying the load costed as much energy as carrying a unit of body mass; 3) as cockroaches adjust to imposed loads, they will become more economical, i.e., cost of carrying loads will decrease. Cockroaches are good models to test these hypotheses because their energetic cost of locomotion and locomotor performance are comparable to other animals (Full, 1989). Additionally, two studies (Lee, 1994; Full et al., 1984, cited in Kram, 1996) suggest they may carry loads economically. From a practical standpoint, cockroaches walk reliably on a treadmill without prior training, even when laden, and they can withstand significant mechanical perturbation without suffering cuticular damage (Jayaram & Full, 2016), making them amenable to artificial loading.

## Materials and Methods

### Animals

Mixed sex, last-instar nymphs of discoid cockroaches (*B. discoidalis*) were purchased from Rainbow Mealworms (Compton, California, USA) in August 2023 and maintained at 26°C, 50% relative humidity and a12:12 light-dark cycle. The colony was provided with puppy and/or kitten chow and water crystals *ad libitum*. Experimental animals were transferred to separate containers and kept under the same conditions.

### Experimental loading

A rectangular piece of lead sheet measuring ∼15 × 20 × 1 mm, depending on cockroach body mass, was affixed dorsally to experimental animals approximately centered above the center of mass (Kram et al.,1997), using self-adhesive strips (Velcro USA Inc., Manchester, NH, USA). In all cases, the combined mass of lead sheet and adhesive strips equaled cockroach body mass on day 0 (see *Experimental protocol* below).

### Respirometry

A respirometry chamber (∼120 × 40 × 60 mm) was made using clear plexiglass and positioned atop a custom-built motorized treadmill (belt width: 50 mm). The respirometry chamber and treadmill were enclosed within a larger container (TICONN Waterproof Electrical Junction Box, inside dimensions: 277 × 176 × 109 mm) so that the lid of this outer container created an airtight ceiling to the inner respirometry chamber when closed (Fig. 1). This outer container had 1 inlet and 3 outlets. The inlet was used to push CO_2_-free air into the container at 5 L × min^-1^. At this rate, it took approximately 8 min to flush the container of ambient air to approximately 0 ppm CO_2_. Narrow slits between the walls of the respirometry chamber and the treadmill belt allowed air to flow from the outer container into the respirometry chamber. The slight positive pressure within the container and flow created by the subsampling pump (described below) prevented air flow in the opposite direction. This was confirmed by tracking the path of injected artificial (vegetable glycerin and propylene glycol) fog into the respirometry chamber. Two small outlets were used to subsample air from both the interior of the outer container and the inner respirometry chamber (Fig. 1). The third larger outlet (∼1 cm diameter) allowed excess CO_2_-free air to flow out of the outer container. Dry CO_2_-free air was generated by a Whatman 75-62 FT-IR purge air generator system (Whatman Inc., Haverhill, MA, USA). Additional scrubbing of this air was performed with ascarite (Arthur H. Thomas Company, Chadds Ford, PA, USA) and drierite columns (W. A. Hammond Drierite Company Ltd., Xenia, OH, USA). Experiments were performed inside an environmental chamber (model I-36VL; Percival Scientific Inc., Boone, IA, USA) from which the door was removed and replaced with clear vinyl sheeting. This allowed easy observation and access while preventing fluctuations in environmental conditions. The temperature inside the environmental chamber was regulated at 26°C, with light intensity set to 370 lux.

**Figure 1.**
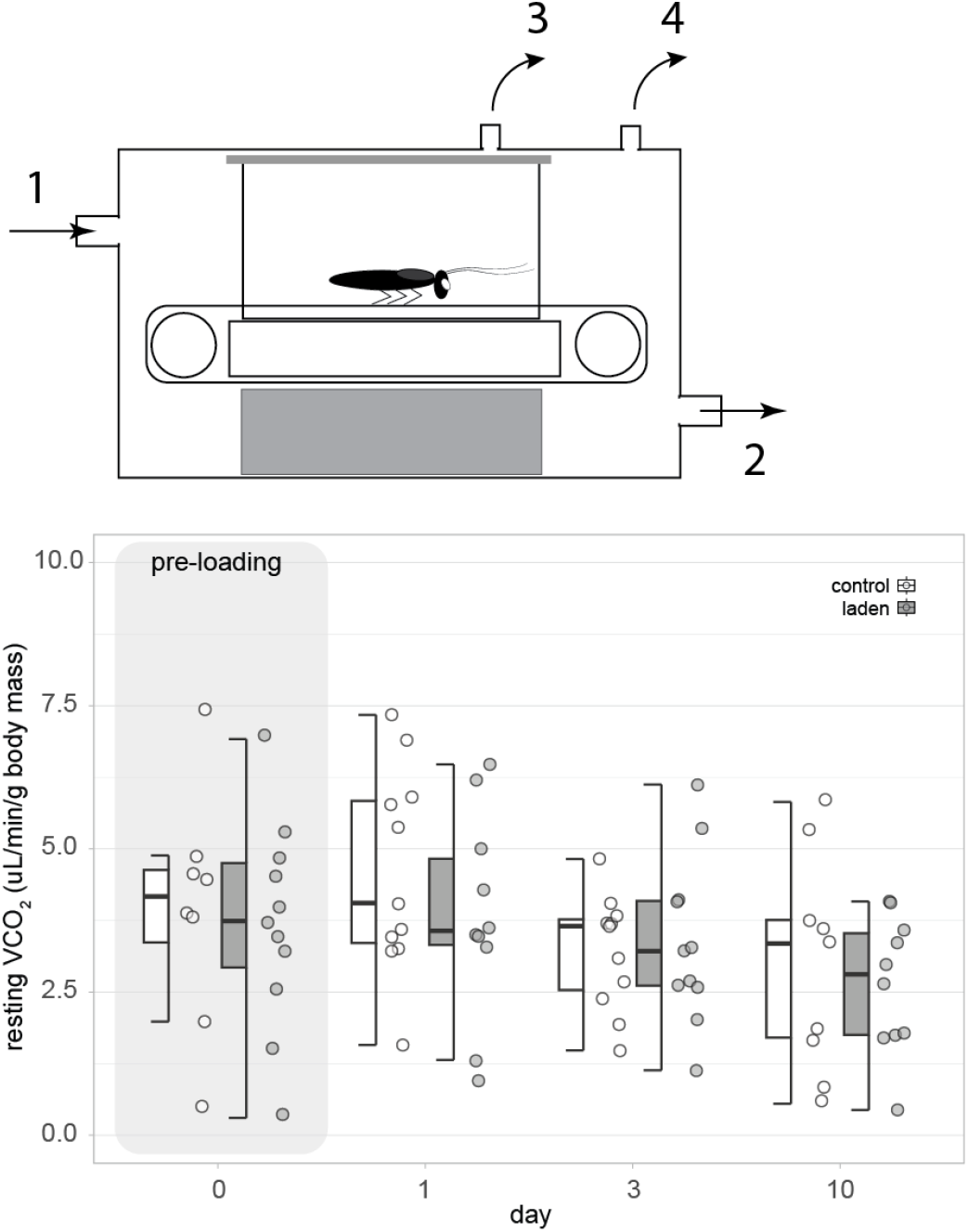
Top panel: Respirometry set-up. The respirometry chamber, placed atop a motorized treadmill, was enclosed within a larger outer container, whose lid created a ceiling to the inner respirometry chamber. Dry CO2-free air was pushed into the container through inlet 1 and subsamples were drawn from the respirometry chamber and the outer container through outlets 3 and 4 respectively. Excess air flowed out of the container through outlet 2. The difference between the CO_2_ concentration measured in subsamples obtained from the respirometry chamber and from the outer container was used to calculate cockroach rate of CO_2_ exchange. Bottom panel: Mass-specific CO_2_ exchange rate during rest (resting VCO_2_) in cockroaches assigned to the laden and control groups on days 0 (pre-loading) and days 1, 3 and 10 after loading onset

Cockroach CO_2_ exchange rates we determined by measuring the difference in CO_2_ concentration between subsampled air from the container and the respirometry chamber, using a LI-COR 7000 CO_2_ sensor system (LI-COR Biosciences, Lincoln, NE, USA). Subsampling flow rates were set to 0.3 L × min^-1^ using two pumps (SS-4 subsampler and the pump of a FoxBox Respirometry system, both from Sable Systems International, North Las Vegas, NV, USA), each connected to a Brooks MFC 5850E mass flow control unit (Costal Instruments, Burgaw, NC, USA) downstream. At this subsampling flow rate, CO_2_ concentration returned to approximately 0 (<4) ppm in less than 2 min following injection of a 30-mL bolus of ambient air directly into the respirometry chamber. Raw LI-COR 7000 data (i.e., air CO_2_ concentration in ppm) was collected by a UI2 data acquisition system controlled by ExpeData software (Sable Systems International, North Las Vegas, NV, USA).

### Experimental protocol

Each animal was weighed and placed inside the respirometry chamber, with the environmental chamber lighting turned off. After a 15-min acclimation period, resting CO_2_ exchange rate was recorded for 5 minutes (resting trial). Aside from minor postural adjustments, most animals remained motionless within a few minutes of being placed in the chamber. Subsequently, lighting was turned on and the treadmill was activated at a low speed (≤0.86 m × min^-1^) for a few minutes to allow animals to adjust to the moving belt. Then, cockroaches completed 4 active trials at progressively faster belt speeds (0.86, 1.40, 1.78, and 2.96 m × min^-1^), for 5 min at each belt speed. This belt speed interval was determined based on pilot experiments to range from the slowest speeds at which animals would move in a straight path, without wandering around the chamber, to the fastest speed that most laden cockroaches could sustain consistently for 5 minutes. Across the range of belt speeds we utilized, cockroaches likely produce energy aerobically (Herreid & Full, 1984), so we assumed that CO_2_ exchange rates directly reflect rates of energy consumption.

This exercise protocol served to establish a baseline for all individuals included in the experiment prior to loading intervention (day 0). After completing the protocol on day 0, animals assigned to the laden experimental group were removed from the respirometry chamber and a piece of lead was attached dorsally, as described, before returning them to the colony. This weight remained attached continuously throughout the rest of the experiment. Control animals were returned to the colony unaltered. After approximately 24 hours of treatment (day 1), animals were retrieved from the colony and repeated the exercise protocol. This was repeated at day 3 and day 10 after loading onset. After completing the protocol on day 10, animals were sacrificed. To ensure laden animals experienced the imposed load throughout the experimental protocol, all cockroaches were encouraged to walk for 30 minutes at their chosen speed on days when they did not exercise on the treadmill. Animals that underwent a molt suffered injuries during the experimental protocol or whose load detached were removed from the experiment.

### Data analysis

Mean mass-specific rate of CO_2_ exchange at rest (resting VCO_2_) was calculated using data from the entire 5-minute (300 s) resting trials. For active trials, mean mass-specific VCO_2_ (active VCO_2_) was calculated as VCO_2_ over the final 150 s of trials at each speed, minus resting VCO_2_. Occasionally, animals were able to sustain a speed long enough to reach a steady state but not for the entire 5 minutes. The last 40-90 seconds were used in these cases. Data was imported from ExpeData (Sable Systems International, North Las Vegas, NV, USA) to R (v4.4.3, R core Team, 2025) within Rstudio (v2025.5.0.496, Posit team, 2025), using the package SableBase (v1.0.0; Foerster, 2013), where it was converted to µL CO_2_ × g ^-1^ × min^-1^. Statistical analyses were performed in R (v4.4.3, R core Team, 2025). Differences in body mass, body length and resting VCO_2_ between control and laden groups on day 0 were tested using two-tailed t-tests with equal variances, as determined by an F-test. The effects of loading on resting VCO_2_ (on days 1-10), and active VCO_2_ (on days 0 and 1-10) were tested using full factorial linear mixed models with restricted maximum likelihood implemented using the lme4 package (v1.1-37., Bates et al., 2015), with treatment, belt speed (for active data) and day (for days 1-10) as main effects and individual as a random effect. Treatment, day, and speed were modeled as categorical variables. If interaction terms were not significant, they were removed from models. Models were examined using type III ANOVA (when interactions were included) or type II ANOVA (when no interaction terms were included) implemented with the R package car (v.3.3-3; Fox & Weisberg, 2019). We performed Tukey-adjusted post-hoc comparisons (using the R package emmeans v1.11.1; Lenth, 2025) among variable groups (i.e., across days, treatments and/or belt speeds). Test results are reported as: test-statistic (degrees of freedom) = value, p = value. Exact p-values are reported unless p < 0.001, in which case they are reported as p < 0.001. The significance level was set at α < 0.05. Resting trials during which animals were agitated and active trials where they did not locomote consistently and/or failed to sustain the speed enough to reach steady state were treated as missing data. Data are presented as mean ± standard deviation unless otherwise specified.

## Results and discussion

### Pre-treatment assessments

26 cockroaches were included in the laden group and 22 in the control group. Of these, 11 (7 male; 4 female) in the laden group and 11 (4 male; 7 female) in the control group completed the experiment. There was no significant difference in body mass (t(20) = −0.02, p = 0.98) and body length (t(20) = 1.83, p = 0.08) between the two experimental groups on day 0 of the experiment. Similarly, day 0 resting VCO_2_ was not different between the two experimental groups (t(17) = 0.28, p = 0.78; note that day 0 resting VCO_2_ was not determined for 3 individuals from the control group; Fig. 1). Day 0 active VCO_2_ increased with belt speed (χ^2^ (3) = 543.84, p<0.001), but did not differ between experimental groups (χ^2^ (1) = 0.77, p = 0.38; Fig 2).

**Figure 2.**
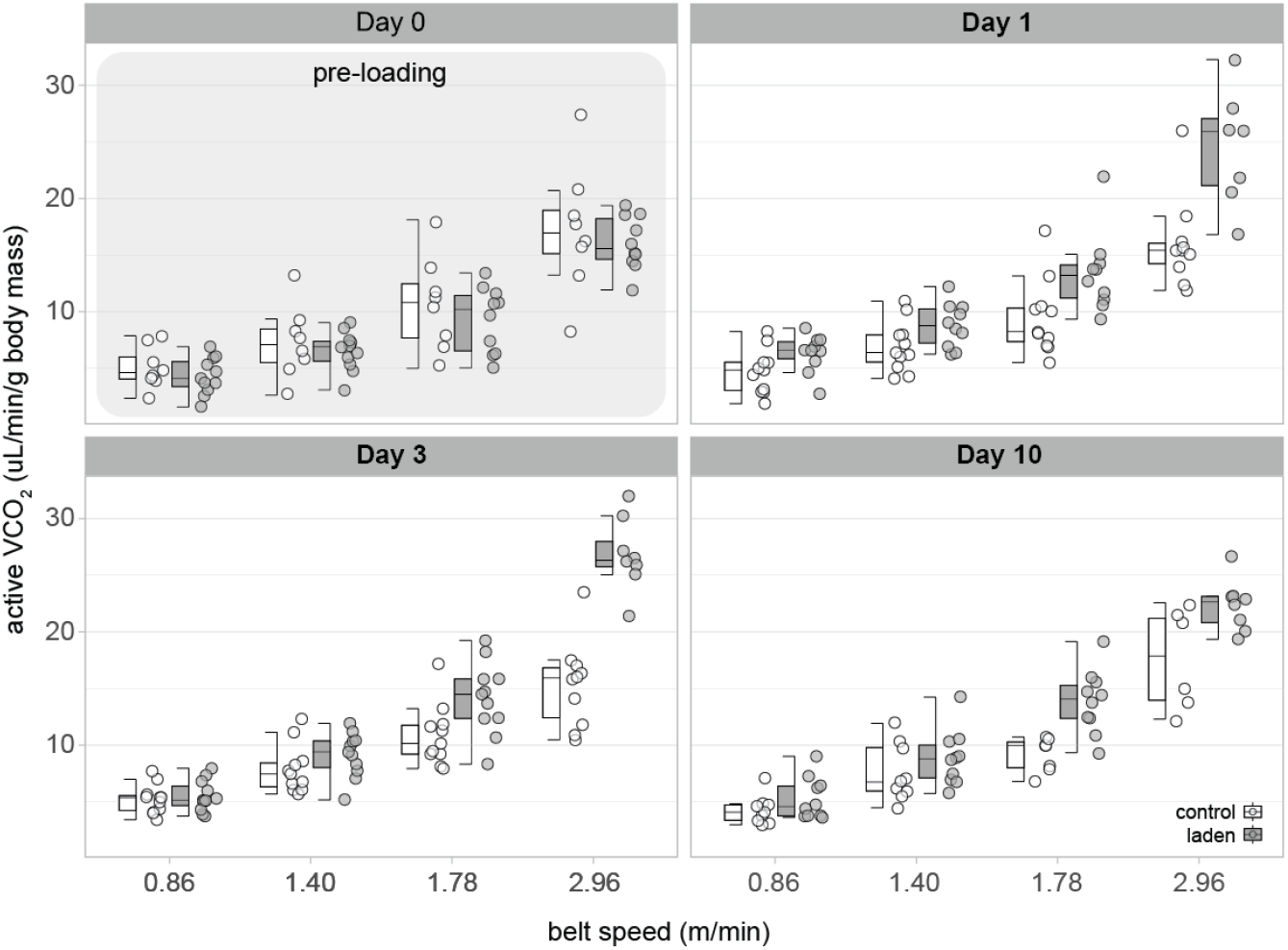
Mass-specific CO_2_ exchange rate during locomotor activity (active VCO_2_) at 4 belt speeds on day 0 (pre-loading), and days 1, 3, and 10 after loading onset.

### Effects of loading on resting VCO_2_

During day 1-10 of the experiment, loading had no significant effect on resting VCO_2_ (χ^2^ (1) = 0.31, p = 0.58; Fig. 1) but it was affected by day of treatment (χ^2^(2) = 11.24, p = 0.004; Fig. 1). Resting VCO_2_ was significantly lower on day 10 compared to day 1 (t(39.9) = 3.23, p < 0.007), while differences between day 1 and day 3 (t(38.5) = 2.34, p = 0.06) and between day 3 and day 10 (t(39.4) = 1.02, p = 0.57) were not significant. These results support our hypothesis that loading does not significantly affect resting VCO_2_ (hypothesis 1).

In many arthropods and small mammals, passive tension generated by the elastic properties of muscles and tendons, and by skeletal or cuticular elements is sufficient - and exceeds - the force needed to counteract gravity (Hooper et al., 2009). This mechanism enables animals with small limbs to maintain posture independently of muscle activation (Yox et al.,1982; Hooper et al., 2009; Dudek & Full, 2006) and may explain the inexpensive load-carrying we demonstrate here, and others have observed in quiescent arthropods (Lee, 1994; Duncan & Lighton, 1994; Lighton et al.,1993; Kram, 1996).

### Effects of loading on active VCO_2_

Active VCO_2_ (Fig 2) varied significantly with belt speed (χ^2^ (3) = 155.62, p < 0.001). The interactions between belt speed and treatment (χ^2^ (3) = 28.29, p < 0.001) and between belt speed, treatment and day (χ^2^ (6) = 22.09, p = 0.001) were also significant. In contrast, the effect of day (χ^2^ (2) = 0.53, p = 0.77) and its interaction with belt speed (χ^2^ (6) = 8.34, p = 0.21) and with treatment (χ^2^ (2) = 1.14, p = 0.57) were not significant. Active VCO_2_ was higher in laden animals at all belt speeds on days 1-10. However, post-hoc tests showed that the difference between treatment groups was not significant at the lower speeds, 0.86 and 1.40 m × min^-1^ (t-ratios ranged from –1.65 to –0.25 and p-values from 0.10 and 0.80). At the higher speeds, 1.78 and 2.96 m × min^-1^, active VCO_2_ was significantly higher in laden animals (t-ratios ranged from –9.31 to –2.68, all p < 0.008). These results partially support hypothesis 2: laden cockroaches used more energy during locomotion than controls. However, at low speeds, the increase was not statistically significant.

In vertebrates, the cost of carrying a load is equal to the cost of moving the same amount of body mass. If this trend holds for cockroaches, active VCO_2_ of laden animals, who were carrying the equivalent of their body mass, should be twice as high as that of controls. While the difference between control and laden animals varied across days and with belt speed, active VCO_2_ of laden cockroaches was never twice as high. On average, across all days and belt speeds, active VCO_2_ of laden cockroaches was 1.33 times that of controls (Fig 3), suggesting that they carry loads economically. Economic load-carrying has been reported in arthropods, but the mechanism that allows them to achieve this remains unclear.

**Figure 3.**
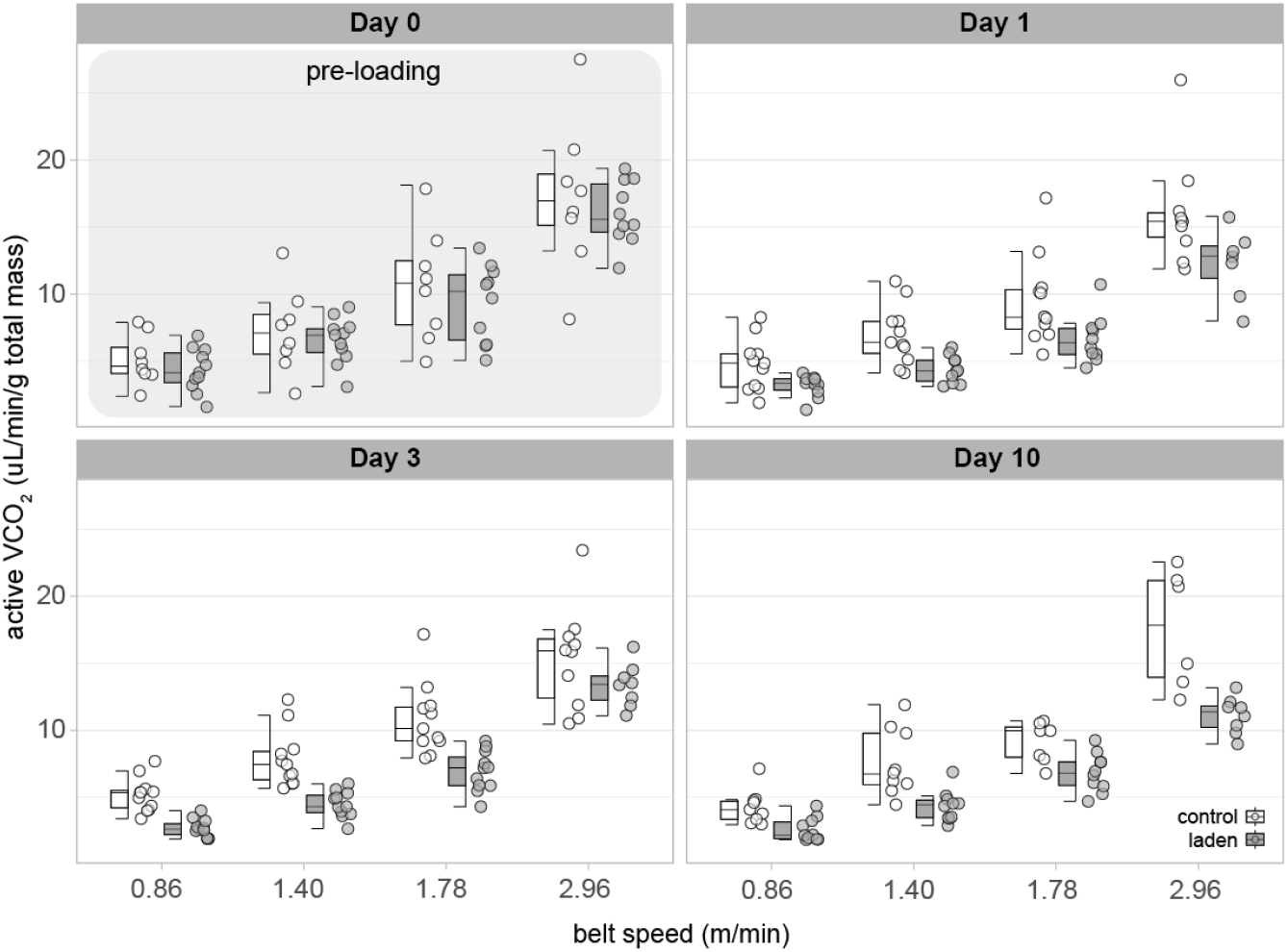
Total mass (body mass plus load mass) – normalized CO_2_ exchange rate during locomotor activity (active VCO_2_) at 4 belt speeds on day 0 (pre-loading), and days 1, 3, and 10 after loading onset.

Nishii (2000) estimated the cost of locomotion in cockroaches, assuming that gait parameters are tuned to minimize this cost. Their results suggest that hexapods may carry loads economically. Our data partially aligns with estimates of that study: cockroaches carry loads economically, but the cost is lower than that predicted by Nishii (2000), who estimates that at a speed of 3 m × min^-1^, carrying the equivalent of their body mass would lead to a ∼90% increase in the cost of locomotion. In the current study, at 2.96 m × min^-1^, active VCO_2_ of laden cockroaches was only ∼50% higher than controls (averaged across days 1, 3 and 10). In addition, Nishii’s (2000) model predicts that cockroaches would carry loads more economically at higher speeds, while our study suggests the opposite relationship. The discrepancy between our results and the predictions of Nishii’s (2000) model may result from its assumption that supporting body and load mass during the stance phase is entirely an active process – i.e., relying on muscle force production. This cost component accounts for a large fraction of the model’s estimated cost of locomotion at low speeds. If skeletal structures aid in supporting the load independently of muscle force, the model’s assumption leads to an overestimation of the energetic cost during absorption phase.

Passive forces may also aid in moving the legs during the propulsion phase. In *Schistocerca gregaria* locusts, forces generated within the femur-tibia joint and by the elastic properties of the extensor tibiae muscle modulate muscle-driven leg extension during cyclic, non-ballistic movements - scratching, walking, and running (Ache & Matheson, 2013). That study shows that the angular velocity of femur-tibia extension is dependent on the initial joint angle from which the leg is released. Release from more flexed femur-tibia joint angles results in faster extensions (Ache & Matheson, 2013). Similar mechanisms aid in joint movements of the false stick insect *Pseudoproscopia scabra* and the stick insect *Carausius morosus* (Ache & Matheson, 2013). Passive force development also aids in the recovery from perturbations in fixed legs of *B. discoidalis* (Dudek & Full, 2006). If passive forces also contribute to movement during walking and running in cockroaches, and if the relative contribution is dependent on the starting angles of leg joints at the start of flexion/extension, changes in kinematics could allow laden animals to take advantage of this mechanism and reduce the energetic cost of locomoting. In Luo and Kikuyu women (Heglund et al., 1995), changes in kinematics allow for an increased energy conservation when carrying loads. Nepalese porters, however, do not rely on this strategy (Bastien et al., 2005; Bastien et al., 2016).

Changes in muscle load, including external loading and manipulations of body mass, lead to adjustments in muscle structure and composition that may impact energetics (e.g., Marden et al., 2001; Marden et al., 2008, Schilder et al. 2011). Indeed, studies that investigated the effects of chronic loading on energetics in mammals (Taylor et al., 1980) and guinea fowl (Katugan-Dechene, 2023) indicate that the energetic cost of carrying loads may decrease over time. Whether this reflects a general pattern among animals remains unknown, particularly for invertebrates.

Contrary to hypothesis 3, we found no systematic change in economy of carrying loads over time: cockroaches carry the equivalent of their body mass economically within 24 h of loading and there is no change in economy in the following days. While changes in leg muscle composition can occur within 24 h in *Drosophila melanogaster* exposed to hypergravity (Schilder & Raynor, 2017), most evidence suggests that adjustments in muscle composition to increases in load take multiple days (Marden et al., 2008; Schilder et al., 2011). Thus, it is not likely that adjustments to muscle composition and functional characteristics facilitate the ability of cockroaches to carry loads economically described here.

We conducted our experiments on last instar *B. discoidalis* nymphs. Kirkton & Harrison (2006) reported changes in jumping performance across ontogenetic development in locusts that align with the ecological demands of each life stage. When forced to jump continuously for 20 minutes, earlier instars, whose survival depends on sustained jumping, develop proportionally less power per jump, but are more resistant to fatigue than later instars and adults. Later instars and adults, who rely on explosive movements, prioritize power development but jump performance quickly declines over time. Therefore, it is possible that the patterns we describe in nymphs of *B. discoidalis* may not be representative of all life stages and cockroach species. In a follow-up study, we will examine the effects of loading on adult energetics in two species of cockroaches with different ecological habits, *B. discoidalis* and *Periplaneta americana*, to shed light on these questions.

Summarizing, in this study we investigate how acute and chronic loading affect the energetics of the discoid cockroach. Our findings indicate that loading has no effect on resting energetics, and that during locomotion, cockroaches carry loads economically. Moreover, the energetic cost of carrying loads does not change as animals adjust to chronic loading. It seems plausible that the ability of these cockroaches to carry significant loads economically may be dependent on changes in locomotor kinematics, but this hypothesis remains to be tested.

## Acknowledgements

We thank Dr. Jaco Klok (Sable Systems) for valuable discussions that were instrumental in designing our respirometer treadmill, Dr. Robert Full for bringing our attention to the energetics of load carrying during rest, and William Lessmann for assisting with cockroach exercise between experimental trials.

## Author contributions

Conceptualization: B.E., R.J.S.; Data curation: B.E., R.J.S; Formal analysis: R.J.S., B. E.; Funding acquisition: R.J.S.; Investigation: B.E., R.J.S.; Methodology: R.J.S., B. E.; Project administration: R.J.S.; Resources: R.J.S, B.E.; Supervision: R.J.S.; Validation: B.E., R.J.S.; Visualization: R.J.S; Writing – original draft: B.E; Writing – review & editing: B.E., R.J.S.

## Data availability

All data and analytical code will be made available from Penn State ScholarSphere (https://scholarsphere.psu.edu/; specific project link TBD).

